# Life-history genotype explains variation in migration activity in Atlantic salmon (*Salmo salar*)

**DOI:** 10.1101/2021.09.14.460042

**Authors:** Petri T. Niemelä, Ines Klemme, Anssi Karvonen, Pekka Hyvärinen, Paul V. Debes, Jaakko Erkinaro, Marion Sinclair-Waters, Victoria L. Pritchard, Laura Härkönen, Craig R. Primmer

**Affiliations:** Organismal and Evolutionary Biology Research Program, Faculty of Biological and Environmental Sciences, University of Helsinki, Helsinki, Finland; University of Jyvaskyla, Department of Biological and Environmental Science, P.O. Box 35, 40014 University of Jyvaskyla, Finland; Natural Resources Institute Finland (Luke), Migratory fish and regulated rivers, Manamansalontie 90, 88300 Paltamo, Finland; Institue of Biotechnology, Helsinki Institute of Life Sciences (HiLIFE), University of Helsinki, Finland; Department of Aquaculture & Fish Biology, Hólar University, Háeyri 1, 551 Sauðárkrókur, Iceland; Natural Resources Institute Finland (Luke), Migratory fish and regulated rivers, Paavo Havaksen tie 3, 90570 Oulu, Finland; Rivers and Lochs Institute, Inverness College, University of the Highlands and Islands, United Kingdom

## Abstract

One of the most important life-history continuums is the fast–slow axis, where “fast” individuals mature earlier than “slow” individuals. “Fast” individuals are predicted to be more active than “slow” individuals; high activity is required to maintain a fast life-history strategy. Recent meta-analyses revealed mixed evidence for such integration. Here, we test whether known life-history genotypes differ in activity expression by using Atlantic salmon (*Salmo salar*) as a model. In salmon, variation in Vgll3, a transcription co-factor, explains ∼40% of variation in maturation timing. We predicted that the allele related to early maturation (*vgll3*\*E) would be associated with increased activity. We used an automated surveillance system to follow ∼1900 juveniles including both migrants and non-migrants (i.e. smolt and parr fish, respectively) in semi-natural conditions over 31 days (∼580 000 activity measurements). Against our prediction, *vgll3* did not explain variation in activity in pooled migrant and non-migrant data. However, in migrants, *vgll3* explained variation in activity according to our prediction in a sex-dependent manner. Specifically, in females the *vgll3*\*E allele was related to increasing activity, whereas in males the *vgll3*\*L allele (later maturation allele) was related to increasing activity. These sex-dependent effects might be a mechanism maintaining within-population genetic life-history variation.

## Introduction

Life-history trade-offs characterize the lives of nearly every organism and when individuals solve trade-offs differently, different life history strategies emerge (Stearns 1992). One of the most studied life-history strategy continuums is the fast–slow continuum, where “fast” life-history strategies have faster developmental rates, younger maturation age and shorter lifespan compared to “slow” strategies with opposite expression of life-history (Stearns 1992; Healy et al. 2019). The fast-slow continuum is one of the most important life-history axes explaining as much as 70% of life-history variation among animal species (Healy et al. 2019). For the last decade, the fast-slow continuum has also been studied at the within-species and within-population levels, with a goal to understand whether and why such continuums exist and how behaviours are associated with them (Réale et al. 2010; Dammhahn et al. 2018a). So far, within-population studies of fast-slow continuums have focused on estimating trait correlation mainly at the un-partitioned phenotypic, but also increasingly at the among-individual, level of (co)variation (Royauté et al. 2018; Moiron et al. 2020). However, to advance our evolutionary understanding of the fast-slow continuum, studies where the genetic underpinning of life-history strategies is known are needed. This would shed more light on the genetic versus environmental contribution on the expression of the continuum and, exclude the need to directly observe life-history events that occur in stages that are difficult or time consuming to track.

Recently, the role of behaviours in expression of fast-slow life-history continuum at the within-population level has received considerable attention (Réale et al. 2010; Dammhahn et al. 2018a; Royauté et al. 2018; Moiron et al. 2020). Behavioural expression is assumed to coevolve with life-history and is considered as a mechanism mediating costs and benefits involved in life-history trade-offs (Stamps 2007; Dammhahn et al. 2018a; Montiglio et al. 2018). For example, individuals maturing at an early age are predicted to be highly explorative and active in order to maintain their “fast” life-history strategy, but might be more vulnerable to predation due to their “risky” behavioural expression. Nevertheless, the results from studies associating behavioural and life-history strategies at the within-population level are mixed (Royauté et al. 2018), possibly because the presence of alternative life-history strategies (e.g. fast, slow) or trade-offs are assumed *a priori* in the majority of empirical studies (Dammhahn et al. 2018). Studying behavioural expression across confirmed life-history genotypes can advance our understanding of whether behaviours and life-history are integrated as suggested by theory.

Salmonid fishes have been a model species group for life-history research (Hendry and Stearns 2004) and this research has contributed substantially to our understanding of the evolution and genetic underpinnings of life-history strategies. For example, in Atlantic salmon (*Salmo salar*), variation in a genome region including the vestigial-like family member 3 gene (*vgll3*) on chromosome 25 has been shown to explain a large proportion of variation in the sea-age at maturity in males and females (Barson et al. 2015). Since *vgll3* is involved in fat cell regulation (Halperin et al. 2013), potentially controlling resource allocation between energy reserves and somatic growth, and its effects are at least partially phylogenetically conserved, it potentially contributes to moderating the expression of life-history strategies across species (Debes et al. 2021). Indeed, *vgll3* is associated with size, as well as age, at maturity in both salmon and humans (Cousminer et al. 2013; Barson et al. 2015). In Atlantic salmon, *vgll3* explains ∼40% of the variation in sea age at maturation so that the *vgll3*\*E allele is linked with early, and the *vgll3*\*L allele with late maturation (Barson et al. 2015). Moreover, early maturing individuals are smaller at maturation compared to late maturing individuals (Barson et al. 2015; Reed et al. 2019). Generally, the life-history of salmon is extremely complex. For example, Atlantic salmon goes through a major life-history transition from the non-migrant (i.e. freshwater *parr* stage) into the migrant life-history stage (i.e. marine *smolt* stage), where physiology, morphology and behavioural expression change dramatically, preparing individuals for transition to the marine environment (Stradmeyer and Thorpe 1987; Huntingford et al. 1988; Thorstad et al. 2012). Altogether, different freshwater and marine life-history stages can last for several years (1-8 & 0-5 years, respectively) (Klemetsen et al. 2003; Mobley et al. 2021), with over 100 different life-history strategies observed in a single population complex (Erkinaro et al. 2019).

Even though many aspects of the association between *vgll3* and life-history are still unknown (e.g. life-history stage-dependent *vgll3* effects on trait expression; difference between migrant and non-migrant), the broad patterns discussed above suggest that generally the *vgll3*\*E allele is linked to “fast” and the *vgll3*\*L allele to “slow” life-histories. This leads to the prediction that the *vgll3*\*E allele is linked to higher activity compared to the *vgll3*\*L allele (Réale et al. 2010; Dammhahn et al. 2018a). So far, there is only indirect evidence that suggest *vgll3* genotypes to differ in their behavioural phenotypes. For example, at sea, *vgll3* homozygote genotypes associate marginally with different prey content in the stomach (Aykanat et al. 2020), which might indicate differences in behaviours contributing to resource acquisition. However, direct behavioural observations are required to fully understand whether behaviours, such as general activity, differ across life-history genotypes or, whether the associations are life-history stage specific.

Using juvenile Atlantic salmon as a model with *vgll3* genotype as a predictor of the life-history strategy, our main objective was to test whether the *vgll3*\*E allele is linked with increased activity. We predicted that the *vgll3*\*E allele would be associated with increased activity in both migrant and non-migrant fish (see above). We measured general movement activity behaviour of 1932 *vgll3*-genotyped, migrant and non-migrant, fish for 31 days using an automatic RFID-surveillance system (i.e. radio frequency identification). Our semi-natural setting (artificial streams) with natural food, water flow and water temperature connects the recorded activity with an ecologically relevant function: activity most likely represents general local, or even territorial, activity in non-migrant fish and migration activity in migrant fish (Stradmeyer and Thorpe 1987; Huntingford et al. 1988; Thorstad et al. 2012). Thus, our study links genetically defined life histories with ecologically relevant behaviours.

## Methods

### Origin of fish

The parents of the experimental fish originated from two broodstocks (Oulujoki; OUL and Tornionjoki; TOR, broodstocks: generation 0), that were hatchery raised at the Natural Resources Institute Finland (Luke), Taivalkoski, but whose parents had successfully completed a sea migration in the Baltic Sea. The breeding design involved crossing only *vgll3* heterozygote individuals, resulting in all possible *vgll3* genotypes occurring within a family in a Mendelian 1:2:1 ratio, thus controlling for genetic background when estimating *vgll3* effects. A series of 2 × 2 factorial crosses were created, i.e. “crossing groups”, using two sires and two dams (one from each parental river) of the *vgll3*\*EL genotype so that both males were crossed with both females. This generated four crossing group types: OUL-OUL, TOR-TOR, TOR-OUL, OUL-TOR with 25 families of each crossing group type. Fish were experimentally crossed in October 2017 and eggs incubated at the Taivalkoski hatchery until hatching, after which eyed eggs (8000 per crossing group, 32000 in total) were transferred to a commercial fish hatchery (Montan Lohi Ltd), where they were maintained for one year in eight fiberglass tanks (3.14 m^2^) (two replicate tanks per crossing group) in similar conditions until February 2019. Because of limited tank capacity, fish from two replicate tanks were combined in one tank on 20^th^ February 2019, leading to one tank per crossing group. In total 3200 fish (800 per crossing group) were tagged between 20-25^th^ February 2019. Fish were anaesthetized (using Benzocaine, 40 mg L^−1^) and a 12 mm passive integrated transponder (HDX PIT-tag, Oregon RFID, Oregon, USA) inserted into the body cavity to enable re-identification and activity recordings. A fin clip was also taken to allow single nucleotide polymorphism (SNP) genotyping. DNA was extracted from fin samples and 177 SNPs including the *VGLL3*TOP SNP (Aykanat et al. 2016) were genotyped and the data subsequently used to assign individuals to families as outlined in Debes et al. (2020). On 27^th^ March 2019 fish were transferred to the Kainuu Fisheries Research Station (www.kfrs.fi, Paltamo, Finland) of the Natural Resources Institute Finland where they were randomly allocated and kept in two fiberglass tanks (15 m^2^) until 23th May 2019 when the experiment was started.

#### Experimental streams and data collection

The experimental streams were circular (surface area = 40m^2^, length = ∼26.15m in the middle, width = 1.5m & depth ∼0.3m) with gravity driven water flow (40.5 L s^-1^, appr. 0.09m s^-1^). Water originated from a nearby lake (Lake Kivesjärvi) mixed from three to seven-meter depth. Thus, the stream water temperature, and food supply (i.e. benthic invertebrates), followed natural variation (Rodewald et al. 2011). The bottom of the stream was covered with gravel (30–80 mm grain size).

Prior to release into the experimental streams, 1932 experimental fish were selected from a larger pool (3200 fish, see above) of PIT tagged and genotyped individuals to ensure an equal representation of *vgll3* genotypes, crossing groups, families and sexes among families. These individuals were then distributed equally among 16 experimental streams (118-124 fish/stream) and left to acclimatize between May 23 – Jun 3, 2019. During the data collection period (31 days; Jun 4 - Jul 4, 2019), fish activity was recorded in the streams. The selected timing of the experiment overlaps with the natural smolt migration timing in Atlantic salmon (Karppinen et al. 2014; Otero et al. 2014). Each stream was equipped with four RFID-antennas, positioned in quadrats of the circular stream (Alioravainen et al. 2020). Since the system had the capacity to record data from maximum 32 antennas at a time, streams were monitored periodically in two sets of eight streams: recording was swapped between the two sets of streams every three days until the end of the data collection period. Thus, each individual stream was monitored on average for 357.4 hours across the 31-day period. The raw-RFID data (theoretical reading frequency 9 readings s^-1^) was converted into one-hour resolution (www.pitdata.net). An individual was defined to have moved if it crossed at least two RFID antennae within the focal hour, i.e. moved through an area between two antennas. As the antennas were equally divided in the stream, the movement measured between two antennas was 6.54 m (one whole round 26.15 m). If an individual did not move during the focal hour, it got value zero (0) for the focal hour movement. We extracted three activity variables from the data. The hourly movement activity was calculated as the number of passes between two antennae in either 1) upstream direction (upstream movement) or 2) downstream direction (downstream movement) and the number of passes between two antennae 3) irrespective of the direction (total movement).

During the observation period, 288 fish disappeared. Either they were missing at the end of the experiment (273 fish) or were observed dead (15 fish) (per genotype: missing; EE=91, EL=87, LL=95, dead; EE=5, EL=7, LL=3). As this resulted in incomplete RFID data from these fish as well as from additional 21 fish that most likely lost their RIFD-tags during the experiment, these fish were removed from the RFID-data. This resulted in a final sample size of 1625 fish (580893 hourly observations of movement), including 500 migrants (178494 hourly observations) and 1125 non-migrants (402399 hourly observations). The migrant status was determined based on a combination of 1) colouration of the fish at the end of the observation period and 2) their movement patterns during the observation period. The transformation of salmon into the migrant saltwater phenotype (smolt) is typically associated with an appearance of silvery colouration and orientation to downstream movement (Thorstad et al. 2012; Debes et al. 2020; Mobley et al. 2021). Here, individuals with a silvery colouration were classified as migrants. Moreover, non-silvery individuals with average downstream movement of at least 5 stream rounds hour^-1^ (i.e. 120 rounds 24h^-1^) were considered as migrant phenotypes (Klemme et al. unpublished). It is good to note that our migrant category includes fish that smoltified (versus the parr that did not) during their first spring (31% of the experimental fish smoltified). Most of the parr were likely to smoltify in later years. At the end of the observation period, fish were used in other experiments.

#### Statistical methods

Generalized linear mixed effect models were used to study whether the number of *vgll3*\*E alleles linked with increased hourly movement activity. Upstream movement, downstream movement and total movement (see above) were each fitted as a response variable in three separate univariate models. Migrant and non-migrant fish are generally known to differ in their behavioural expression and the biological meaning of the recorded activity likely differs between individuals with different migrant status (e.g. migration activity versus local activity) (Stradmeyer and Thorpe 1987; Huntingford et al. 1988; Thorstad et al. 2012). Thus, we ran separate univariate models for i) data where migrant and non-migrant fish were pooled, ii) only migrant fish and iii) only non-migrant fish for each three recorded activity.

The main models were fitted with *vgll3* having additive (i.e. EE = 1, EL = 0 and LL = -1; continuous covariate) and dominance (i.e. EE & LL = 0 and EL = 1; continuous covariate) effects on activity (Xiang et al. 2018). Fitting both additive and dominance *vgll3* effects in the model estimates whether there are dominance effects on top of additive effects (Xiang et al. 2018). Additionally, fixed effects for sex (categorical) and migrant status (migrant or non-migrant, categorical) were fitted with *vgll3* interaction. Interaction terms between *vgll3* and sex were considered since in some cases, the *vgll3*\*E allele has been observed to be dominant in males, but not in females (Barson et al. 2015). Moreover, interactions with migrant status were considered as the behavioural interpretation of the measured activity likely differs between smolt and parr (Stradmeyer and Thorpe 1987; Huntingford et al. 1988; Thorstad et al. 2012). Migrant status was fitted only in models where pooled data were used. All main models can be found in Table 1 and Supplementary Table 1.

**Table 1.** Parameter estimates for additive and dominance models for three behaviours: upstream activity, downstream activity and total activity. We present fixed parameter estimates (i.e. β, on the log scale) with standard error (i.e. SE) and *P*-values and, random parameter estimates (i.e. standard deviation = σ) with 95% credible intervals (i.e. CI). Genotypes modeled as: *vgll3*\*EE = 1, *vgll3*\*EL = 0, *vgll3*\*LL = -1.

| **Pooled phenotypes** | *Upstream activity* |  | *Downstream activity* |  | *Total activity* |  |
| --- | --- | --- | --- | --- | --- | --- |
| **Fixed effects** | $\beta$ (SE) | *P* | $\beta$ (SE) | *P* | $\beta$ (SE) | *P* |
| Intercept | -0.958 (0.232) | >0.001 | 0.796 (0.229) | >0.001 | 1.220 (0.219) | >0.001 |
| *vgll3*_ADDITIVE | -0.025 (0.044) | 0.571 | -0.005 (0.043) | 0.909 | -0.006 (0.040) | 0.882 |
| Sex<sup>1</sup> | -0.087 (0.053) | 0.101 | -0.066 (0.052) | 0.207 | -0.068 (0.049) | 0.159 |
| *vgll3*_DOMINANCE | -0.064 (0.074) | 0.381 | -0.043 (0.072) | 0.551 | -0.043 (0.067) | 0.523 |
| Migrant status<sup>2</sup> | 2.085 (0.061) | >0.001 | 2.131 (0.060) | >0.001 | 2.021 (0.056) | >0.001 |
| *vgll3*_ADDITIVE:Sex<sup>1</sup> | 0.042 (0.053) | 0.431 | 0.034 (0.052) | 0.515 | 0.033 (0.048) | 0.489 |
| *vgll3*_DOMINANCE:Sex<sup>1</sup> | 0.042 (0.091) | 0.640 | 0.010 (0.089) | 0.915 | 0.015 (0.083) | 0.855 |
| *vgll3*_ADDITIVE:Migrant status<sup>2</sup> | 0.005 (0.056) | 0.934 | -0.010 (0.056) | 0.856 | -0.009 (0.052) | 0.865 |
| *vgll3*_DOMINANCE:Migrant status<sup>2</sup> | 0.044 (0.097) | 0.651 | 0.059 (0.096) | 0.536 | 0.054 (0.089) | 0.545 |
| **Random Effects** | $\sigma$ (95% CI) | | $\sigma$ (95% CI) | | $\sigma$ (95% CI) | |
| Individual | 0.829 (0.798, 0.861) |  | 0.818 (0.787, 0.849) |  | 0.763 (0.735, 0.791) |  |
| Date | 1.082 (0.843, 1.389) |  | 1.067 (0.831, 1.369) |  | 1.020 (0.795, 1.308) |  |
| Stream | 0.201 (0.131, 0.307) |  | 0.203 (0.134, 0.310) |  | 0.186 (0.122, 0.284) |  |
| Crossing group | 0.145 (0.053, 0.395) |  | 0.140 (0.051, 0.385) |  | 0.132 (0.048, 0.361) |  |
| Mother | 0.162 (0.093, 0.281) |  | 0.153 (0.087, 0.271) |  | 0.145 (0.083, 0.254) |  |
| Father | 0.229 (0.161, 0.326) |  | 0.219 (0.153, 0.312) |  | 0.198 (0.138, 0.285) |  |
| Hour | 0.303 (0.227, 0.404) |  | 0.326 (0.244, 0.434) |  | 0.319 (0.239, 0.425) |  |
| **Migrant fish** | *Upstream* |  | *Downstream* |  | *Total activity* |  |
| **Fixed effects** | $\beta$ (SE) | *P* | $\beta$ (SE) | *P* | $\beta$ (SE) | *P* |
| Intercept | 0.658 (0.309) | 0.033 | 2.448 (0.317) | >0.001 | 2.774 (0.306) | >0.001 |
| *vgll3*_ADDITIVE | 0.049 (0.029) | 0.097 | 0.065 (0.036) | 0.073 | 0.064 (0.036) | 0.072 |
| Sex<sup>1</sup> | -0.038 (0.039) | 0.335 | -0.022 (0.049) | 0.649 | -0.025 (0.048) | 0.595 |
| *vgll3*_DOMINANCE | 0.082 (0.048) | 0.087 | 0.111 (0.059) | 0.063 | 0.109 (0.059) | 0.064 |
| *vgll3*_ADDITIVE:Sex<sup>1</sup> | -0.108 (0.039) | 0.006 | -0.131 (0.048) | 0.007 | -0.131 (0.047) | 0.006 |
| *vgll3*_DOMINANCE:Sex<sup>1</sup> | -0.099 (0.068) | 0.143 | -0.127 (0.085) | 0.134 | -0.130 (0.166) | 0.118 |
| **Random Effects** | $\sigma$ (95% CI) | | $\sigma$ (95% CI) | | $\sigma$ (95% CI) | |
| Individual | 0.340 (0.316, 0.365) |  | 0.424 (0.395, 0.454) |  | 0.416 (0.388, 0.446) |  |
| Date | 1.566 (1.221, 2.009) |  | 1.574 (1.227, 2.019) |  | 1.511 (1.177, 1.938) |  |
| Stream | 0.308 (0.210, 0.452) |  | 0.291 (0.193, 0.437) |  | 0.280 (0.186, 0.421) |  |
| Crossing group | 0.165 (0.069, 0.394) |  | 0.201 (0.084, 0.481) |  | 0.200 (0.084, 0.476) |  |
| Mother | 0.051 (0.014, 0.178) |  | 0.055 (0.012, 0.252) |  | 0.055 (0.012, 0.244) |  |
| Father | 0.072 (0.026, 0.203) |  | 0.106 (0.047, 0.238) |  | 0.102 (0.045, 0.235) |  |
| Hour | 0.212 (0.158, 0.283) |  | 0.266 (0.199, 0.355) |  | 0.259 (0.194, 0.346) |  |
| **Non-migrant fish** | *Upstream activity* |  | *Downstream activity* |  | *Total activity* |  |
| **Fixed effects** | $\beta$ (SE) | *P* | $\beta$ (SE) | *P* | $\beta$ (SE) | *P* |
| Intercept | -0.856 (0.237) | >0.001 | 0.642 (0.231) | 0.006 | 1.106 (0.216) | >0.001 |
| *vgll3*_ADDITIVE | -0.061 (0.071) | 0.387 | -0.047 (0.069) | 0.496 | -0.048 (0.064) | 0.451 |
| Sex<sup>1</sup> | -0.141 (0.095) | 0.136 | -0.123 (0.093) | 0.186 | -0.127 (0.086) | 0.141 |
| *vgll3*_DOMINANCE | -0.094 (0.0117) | 0.418 | -0.071 (0.0115) | 0.537 | -0.070 (0.106) | 0.508 |
| *vgll3*_ADDITIVE:Sex<sup>1</sup> | 0.091 (0.094) | 0.334 | 0.097 (0.092) | 0.293 | 0.098 (0.085) | 0.248 |
| *vgll3*_DOMINANCE:Sex<sup>1</sup> | 0.099 (0.159) | 0.533 | 0.060 (0.156) | 0.703 | 0.073 (0.144) | 0.612 |
| **Random Effects** | $\sigma$ (95% CI) | | $\sigma$ (95% CI) | | $\sigma$ (95% CI) | |
| Individual | 1.028 (0.974, 1.085) |  | 1.013 (0.960, 1.068) |  | 0.936 (0.888, 0.987) |  |
| Date | 0.822 (0.640, 1.056) |  | 0.785 (0.611, 1.009) |  | 0.734 (0.571, 0.943) |  |
| Stream | 0.262 (0.159, 0.433) |  | 0.280 (0.172, 0.456) |  | 0.256 (0.157, 0.417) |  |
| Crossing group | 0.177 (0.062, 0.511) |  | 0.170 (0.057, 0.506) |  | 0.158 (0.053, 0.467) |  |
| Mother | 0.289 (0.176, 0.477) |  | 0.290 (0.176, 0.479) |  | 0.269 (0.164, 0.443) |  |
| Father | 0.316 (0.211, 0.471) |  | 0.320 (0.216, 0.474) |  | 0.292 (0.196, 0.434) |  |
| Hour | 0.493 (0.370, 0.657) |  | 0.476 (0.357, 0.635) |  | 0.456 (0.342, 0.608) |  |

<sup>1</sup> Reference sex is female, <sup>2</sup> Reference migrant status is non-migrant

In all models, we included the same random effects structure. We fitted individual identity as a random effect to control for pseudo-replication. To control for variation in the data caused by family structure and potential spatiotemporal variation caused by the experimental setup, we fitted mother identity (34 mothers), father identity (42 fathers), crossing group (i.e. parental source population combination (four levels: OUL-OUL, TOR-TOR, TOR-OUL, OUL-TOR), stream identity (16 streams), date (31 days) and time of day (hour identity; 24 levels) as random effects.

All models were run in the R statistical environment R, version 3.6.3. (R core team 2020), using the package glmmTMB (Brooks et al. 2017) with “negative binomial” error distribution and log-link function, which fits well for zero-inflated count data.

## Results

*Vgll3* was not linked with upstream, downstream or total activity when data pooled across migrant phenotypes were used (Table 1, Figure 1). Moreover, in non-migrant fish, *vgll3* effects were absent.

**Figure 1.**
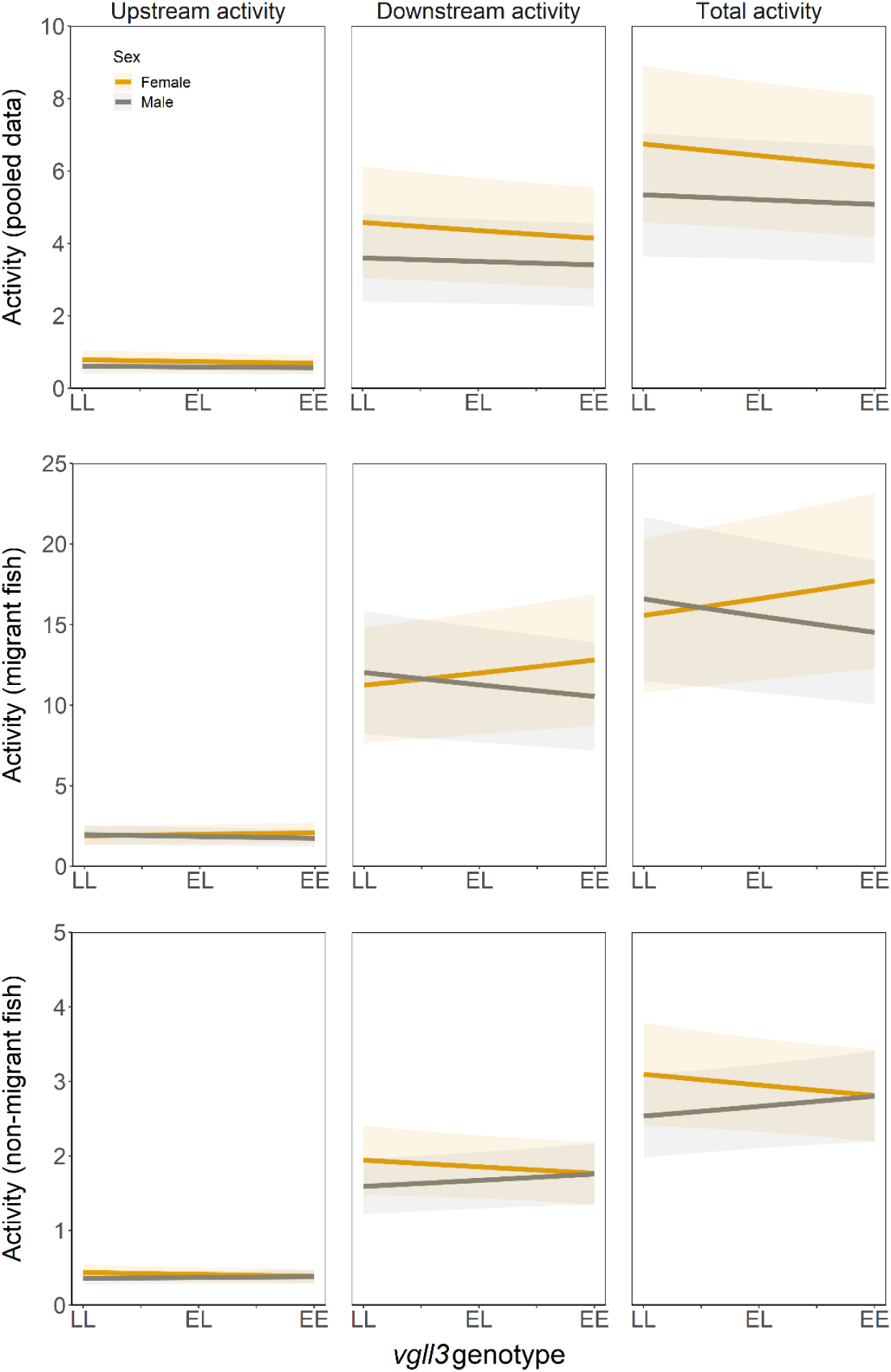
Linear predictions for *vgll3* additive effects for all three behaviours and all three data types (i.e. migrant, non-migrant and pooled data). The predictions are derived from model estimates presented in Table 1. Shaded areas represent standard errors around the predictions.

In migrant fish, *vgll3* effects on upstream, downstream and total activity were sex-dependent (Table 1, Figure 1). More specifically, the *vgll3*\*E allele, which relates to earlier maturation, was linked with increased activity in females. However, in males *vgll3*\*L, which relates to later maturation, was linked with increased activity (Table 1, Figure 1) (see Supplementary Table 2 for data scale predictions, used in Figure 1, and for activity expressed as distance moved in meters).

The dominance effects were non-significant (Table 1), although the effect sizes for additive and dominance effects were quite similar and statistical significance for dominance was close to the threshold value. Thus, our results are inconclusive.

Variation for date effects was significantly larger for migrant and variation for individual effects was marginally larger for non-migrant fish compared to other random effects, when 95% credible intervals across random effects are being compared (Table 1, Supplementary Figure 1). This means that day-to-day differences dominated activity variation in migrant fish (Supplementary Figure 1). To test whether date effects explain the observed *vgll3* effects, we ran an additional model where we included *vgll3-*date interaction in the main model (Supplementary Table 3). Nevertheless, when we added *vgll3*-date interaction as a random effect in the migrant fish model, the *vgll3* effect estimate on migrant activity remained the same (Supplementary Table 3). Finally, among-individual differences in activity dominated activity variation in non-migrant fish (Table 1).

## Discussion

In this study, we tested whether individuals with genetically determined life-history strategies differ in their activity levels as predicted by pace-of-life theory (Réale et al. 2010; Dammhahn et al. 2018b; Laskowski et al. 2021), specifically whether the *vgll3*\*E allele, which associates with earlier maturation, is linked with increased activity. Interestingly, *vgll3* effects on activity differed between males and females within migrant fish and our predictions were supported only in migrant females, where the *vgll3*\*E allele was indeed linked with increased activity. In migrant males, conversely, the *vgll3*\*L allele, which relates to later maturation, was linked with increased activity. In non-migrant fish, *vgll3* effects were absent. Thus, our results indicate that *vgll3* effects on activity depend on migrant status and sex.

### Migrant status and sex-dependent *vgll3* effects on activity

Our results indicate that variation in *vgll3* genotype explains variation in migration activity. Increased hourly migration activity might allow individuals to reach the marine environment considerably faster. Indeed, the literature acknowledges considerable variation in the duration of the (smolt) migration among individuals (Thorstad et al. 2012; Karppinen et al. 2014). Our results indicate that sex-dependent variation in *vgll3* genotype might partly explain this variation so that the *vgll3*\*E allele is linked with increased migration activity in females, while in males the pattern is opposite (the result could also be explained by genotype and sex-specific onset of migration, but no such *vgll3*-sex interaction was observed: Supplementary Table 4). Generally, smolt migration takes several days, up to 8 weeks, depending on the river length (Thorstad et al. 2012). Thus, even small differences in hourly migration activity across genotypes could lead to large differences in the overall migration duration in long rivers. Lower migration activity can be considered as a more “risky” behaviour compared to high migration activity (Thorstad et al. 2012; Hyvärinen and Rodewald 2013; Karppinen et al. 2014). Indeed, fast migration through estuaries or other high-risk areas has been suggested to reduce mortality during migration to the sea (reviewed in Thorstad et al. 2012). For example, multiple fish, bird and mammal species prey on migrating fish in the river (Thorstad et al. 2012; Karppinen et al. 2014). Faster migrating *vgll3* genotypes might be able to (temporarily) reduce migration-related mortality costs, compared to other genotypes, leading to genotype-dependent survival during the river-to-sea migration. Our finding of a sex-genotype interaction for migration activity most likely arises due to sex-dependent expression of life-history (trade-offs) (see below), which affects the costs and benefits of behavioural expression in a sex-dependent manner (Immonen et al. 2018). Interestingly, under the assumption that variation in migration speed generates variation in survival, the sex-*vgll3* interaction in migration activity might act as a mechanism maintaining variation in *vgll3* genotypes: opposite genotypes in males (i.e. LL) and females (i.e. EE) might be selected for due to viability selection acting via migration duration/speed. On the other hand, such antagonistic selection can make sex-specific adaptations, linked to *vgll3* genotypes, inherently difficult.

Potential mechanisms explaining variation in migration activity linked to *vgll3* genotype might be variation in body condition as it might affect behavioural expression in salmon (Thorstad et al. 2012). Furthermore, if the effects of body condition on migration activity are sex-dependent, that might partly explain the patterns found in the current study. Indeed, when testing this *a posteriori* hypothesis, adding an interaction between sex and body condition in our (migrant fish) models rendered the *vgll3* effects on activity much smaller and non-significant (Supplementary Table 1). Amongst individuals in poorer condition, males had higher migration activity than females while amongst individuals with higher condition, we found the opposite pattern (Supplementary figure 2). It has been previously shown that the *vgll3*\*E allele is associated with higher condition (Debes et al. 2021), which was also (although marginally) confirmed by our *a posteriori* analysis in migrant fish (Supplementary Figure 3). Thus, variation in body condition might explain the sex - *vgll3* interaction in migration activity: in females, the *vgll3*\*E allele, i.e. which relates to higher condition, is also associated with increased migration activity while in males, the *vgll3*\*L allele, which relates to lower condition, is associated with higher migration activity. Interestingly, body condition has been suggested to, at least partly, be a factor contributing to the detected *vgll3* effects on maturation timing of male parr (Debes et al. 2021). Our results add more evidence supporting the notion that *vgll3* effects on trait expression might be mediated by body condition, although more research on the topic is needed.

#### Fast-slow life-history continuum and movement activity

Our results indicate that only in migrant females does *vgll3* genotype explain movement activity as predicted by pace-of-life theory (Réale et al. 2007; Dammhahn et al. 2018a). Sex-dependent expression of the pace-of-life continuum, e.g. integration of life-history and behavioural expression, has been predicted before, but rarely studied (Immonen et al. 2018; Tarka et al. 2018). The potential dependence of this life history-behaviour integration on ecology might explain why the *vgll3*-activity association is expressed in opposite ways in males and females (Immonen et al. 2018; Montiglio et al. 2018; Tarka et al. 2018; Laskowski et al. 2021). In Atlantic salmon, males and females differ in the expression of key life-history traits and how these traits affect fitness. For example, males generally mature earlier compared to females (Fleming and Einum 2011; Erkinaro et al. 2019) and females benefit more from larger maturation size compared to males (Fleming and Einum 2011; Mobley et al. 2020). In addition, sexes differ in expression of the trade-off between time spent in freshwater versus marine environments (Mobley et al. 2020). Sex differences in life-history (trade-offs) might lead to males destined to mature late but females destined to mature early to adopt increased migration activity. The current general theory predicting links between life-history and behavioural expression is quite broad and not sufficiently detailed to explain why there are so many exceptions, including our study, to the fast-slow continuum (Immonen et al. 2018; Montiglio et al. 2018; Laskowski et al. 2021).

## Conclusions

Variation in *vgll3* in Atlantic salmon has previously been shown to explain variation in the expression of life-history strategies in males and females (Barson et al. 2015). Here, we show that *vgll3* explains variation in movement activity. Our work reveals complex behaviour – life history integration, as the prediction from the pace-of-life theory was supported only in migrant females, where higher activity rate relates to the genetic predisposition for earlier maturation age. Future empirical and modeling work may consider sex and life-history stage-dependent expression of behaviour-life history integration since the theory, in its current form, does not consider such dependency. This may be one of the reasons why the theory is generally not well supported by empirical studies (Royaute et al. 2018; Moiron et al. 2020; Laskowski et al. 2021).

## Acknowledgements

This work was supported by the Academy of Finland (project numbers 284941, 286334, 314254 and 314255), the European Research Council under the European Articles Union’s Horizon 2020 research and innovation program (grant no. 742312), European Maritime and Fisheries Fund (grant no. 43521) and the University of Helsinki. We also thank the personnel of Montan Lohi Oy fish farm, Taivalkoski fish farm and Kainuu fisheries research station for raising and maintaining the salmon population. We thank Jyrki Oikarinen, Olli van der Meer, Juuso Meriläinen, Mikko Jaukkuri, Ari Leinonen, Kanerva Korhonen, Tapio Laaksonen, Pekka K. Korhonen, Rauno Hokki ja Vesa Rantanen for assisting with marking of the fish and organizing the experimental setup. We thank Andrew House, Annukka Ruokolainen, Jacqueline Moustakas-Verho, Johanna Kurko, Ksenia Zueva, Nico Lorenzen, with their help in generating the crossed families. We also thank Annukka Ruokolainen and Seija Tillanen for conducting the SNP genotyping. We declare no conflict of interest.

## Open data

The data and R-script related to this study are stored in public repository when the manuscript is accepted for publication.

---

**Supplementary Table 1.** Parameter estimates for additive and dominance models for three behaviours when body condition is added as a covariate in the migrant fish model: upstream activity, downstream activity and total activity (i.e. downstream and upstream activity pooled). We present fixed parameter estimates (i.e. β) with standard error (i.e. SE) and *P*-values and, random parameter estimates (i.e. standard deviation = σ) with 95% credible intervals (i.e. CI). Genotypes modeled as: *vgll3*\*EE = 1, *vgll3*\*EL = 0, *vgll3*\*LL = -1.

| **Migrant fish** | *Upstream activity* |  | *Downstream activity* |  | *Total activity* |  |
| --- | --- | --- | --- | --- | --- | --- |
| **Fixed effects** | $\beta$ (SE) | *P* | $\beta$ (SE) | *P* | $\beta$ (SE) | *P* |
| Intercept | 0.704 (0.310) | 0.023 | 2.513 (0.320) | >0.001 | 2.837 (0.309) | >0.001 |
| *vgll3*_ADDITIVE | -0.001 (0.055) | 0.983 | <-0.001 (0.068) | 0.998 | -0.002 (0.067) | 0.978 |
| Sex<sup>1</sup> | -0.071 (0.054) | 0.189 | -0.065 (0.067) | 0.335 | -0.072 (0.066) | 0.277 |
| *vgll3*_DOMINANCE | -0.109 (0.093) | 0.240 | -0.148 (0.115) | 0.199 | -0.143 (0.113) | 0.205 |
| Condition | -0.048 (0.016) | 0.002 | -0.063 (0.020) | 0.001 | -0.063 (0.019) | 0.001 |
| *vgll3*_ADDITIVE:Sex<sup>1</sup> | 0.043 (0.075) | 0.568 | 0.050 (0.093) | 0.594 | 0.047 (0.091) | 0.608 |
| *vgll3*_DOMINANCE:Sex<sup>1</sup> | 0.097 (0.130) | 0.455 | 0.123 (0.162) | 0.449 | 0.129 (0.159) | 0.415 |
| Condition:Sex<sup>1</sup> | -0.047 (0.021) | 0.029 | -0.053 (0.026) | 0.043 | -0.051 (0.026) | 0.048 |
| **Random Effects** | $\sigma$ (95% CI) | | $\sigma$ (95% CI) | | $\sigma$ (95% CI) | |
| Individual | 0.321 (0.299, 0.344) |  | 0.400 (0.373, 0.429) |  | 0.392 (0.366, 0.420) |  |
| Date | 1.566 (1.221, 2.010) |  | 1.574 (1.227, 2.020) |  | 1.511 (1.178, 1.939) |  |
| Stream | 0.306 (0.208, 0.449) |  | 0.298 (0.198, 0.447) |  | 0.284 (0.189, 0.427) |  |
| Crossing group | 0.171 (0.072, 0.403) |  | 0.209 (0.089, 0.489) |  | 0.206 (0.088, 0.482) |  |
| Mother | 0.014 (<0.001, >10.00) |  | 0.002 (<0.001, >10.00) |  | 0.002 (<0.001, >10.00) |  |
| Father | 0.100 (0.052, 0.192) |  | 0.129 (0.074, 0.227) |  | 0.129 (0.074, 0.225) |  |
| Hour | 0.212 (0.158, 0.283) |  | 0.266 (0.199, 0.355) |  | 0.259 (0.194, 0.346) |  |

<sup>1</sup> Reference sex is female

**Supplementary Table 2.**
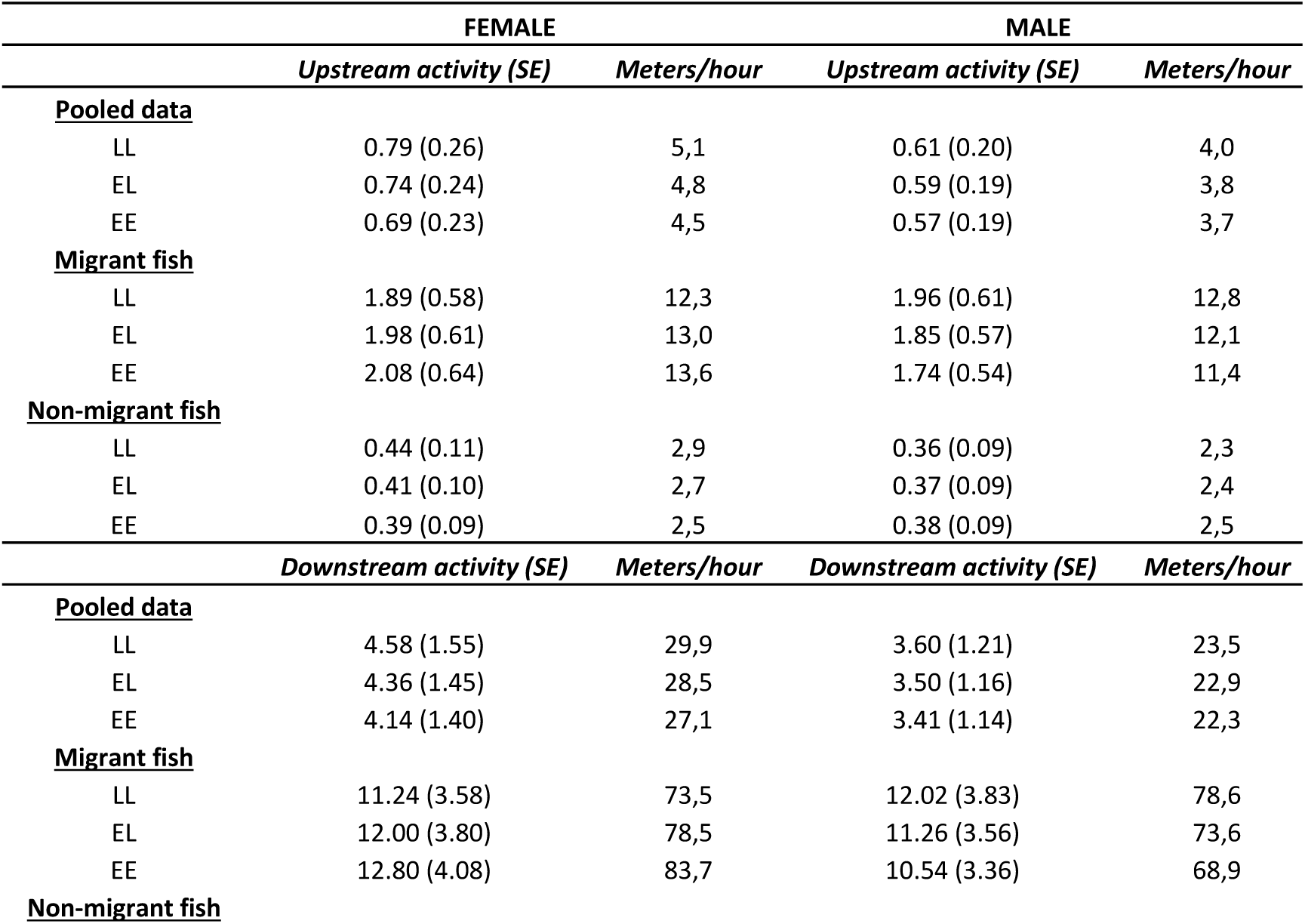
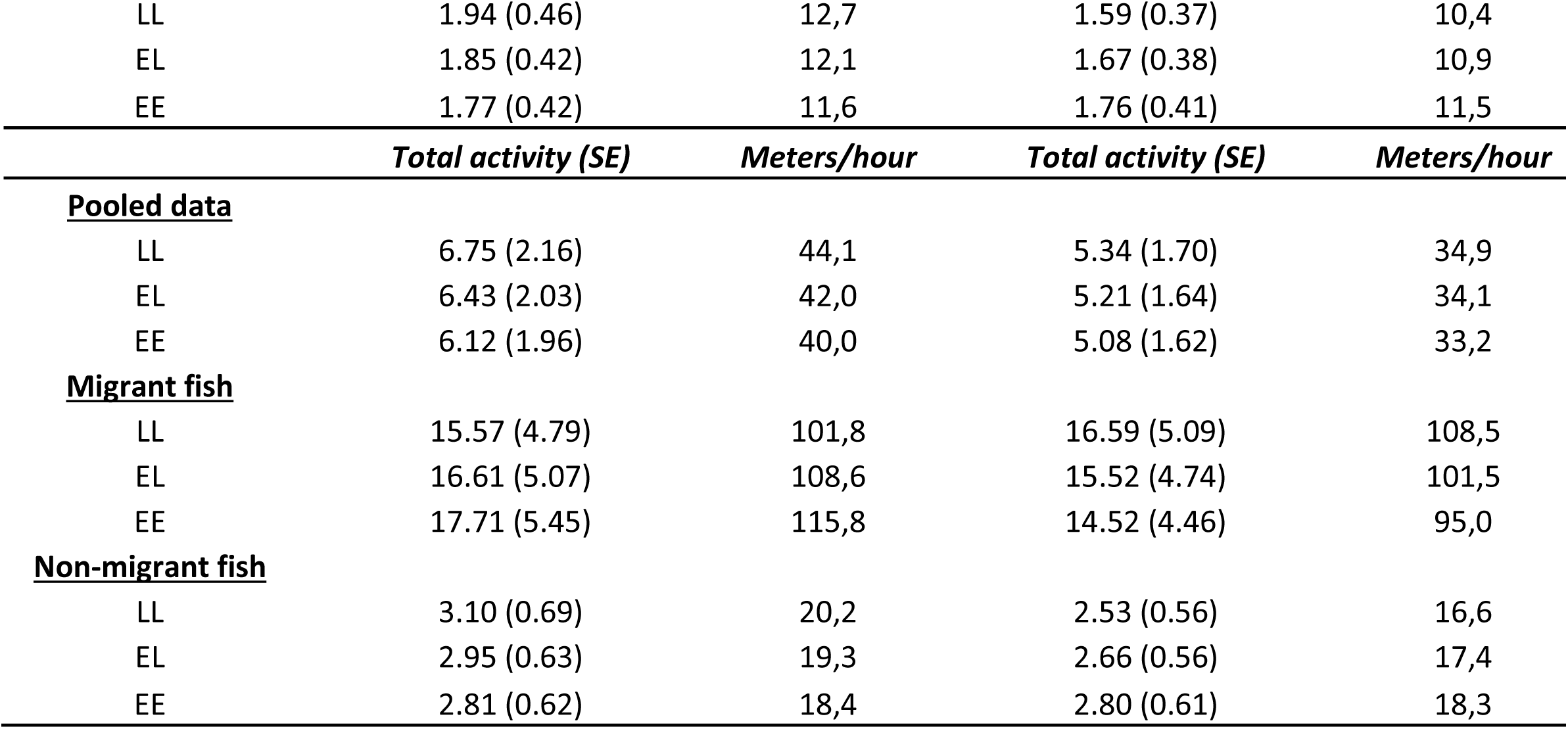
Data scale predictions and hourly distance moved in meters for *vgll3* and sex effects on activity. Predictions are extracted from models presented in Table 1. The conversion to meters was done by multiplying the activity estimates by 6.54, i.e. the distance between two adjacent antennae.

**Supplementary Table 3.** Parameter estimates for a model where a *vgll3*-date interaction is fitted as a random effect for the model presented in Table 1. We present fixed parameter estimates (i.e. β) standard errors (i.e. SE) and P-values. As the results were qualitatively the same across all three behaviours, we present here only the estimates for downstream activity.

| **Migratory fish** | *Downstream activity* |  |
| --- | --- | --- |
| **Fixed effects** | $\beta$ (SE) | *P* |
| Intercept | 2.447 (0.317) | >0.001 |
| *vgll3_ADDITIVE* | 0.087 (0.038) | 0.022 |
| Sex<sup>1</sup> | -0.022 (0.049) | 0.647 |
| *vgll3_DOMINANCE* | 0.111 (0.060) | 0.063 |
| *vgll3_ADDITIVE:Sex*<sup>1</sup> | -0.133 (0.048) | 0.006 |
| *vgll3_DOMINANCE:Sex*<sup>1</sup> | -0.126 (0.085) | 0.136 |
| **Random Effects** | $\sigma$ | *r* |
| Individual | 0.424 |  |
| Date | 1.574 |  |
| *vgll3_ADDITIVE* | 0.054 | -0.61 |
| Stream | 0.290 |  |
| Crossing group | 0.202 |  |
| Mother | 0.054 |  |
| Father | 0.106 |  |
| Hour | 0.266 |  |

<sup>1</sup> Reference sex is Female

**Supplementary Table 4.**
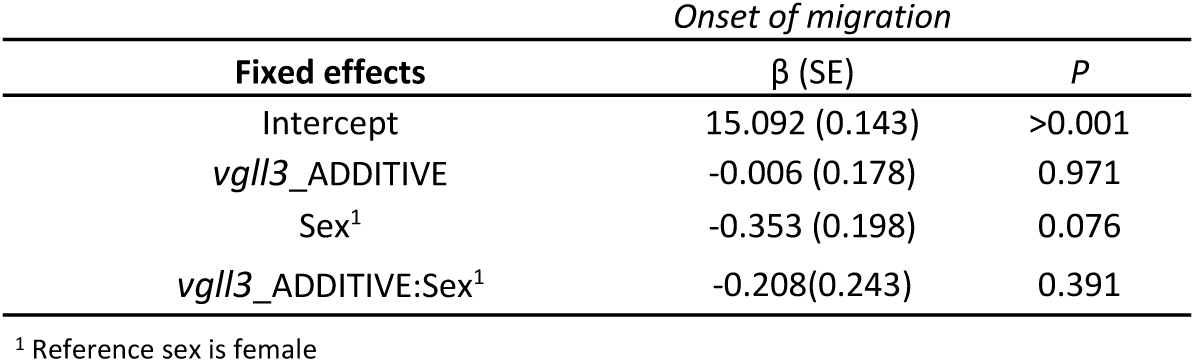
Parameter estimates for *vgll3* and sex effects on onset of migration. We present fixed parameter estimates (i.e. β) standard errors (i.e. SE) and P-values. Onset of migration was defined as date when the downstream movement reached 120 stream rounds day^-1^ (i.e. 5 rounds hour^-1^) among migrant fish (see methods).

**Supplementary Figure 1.**
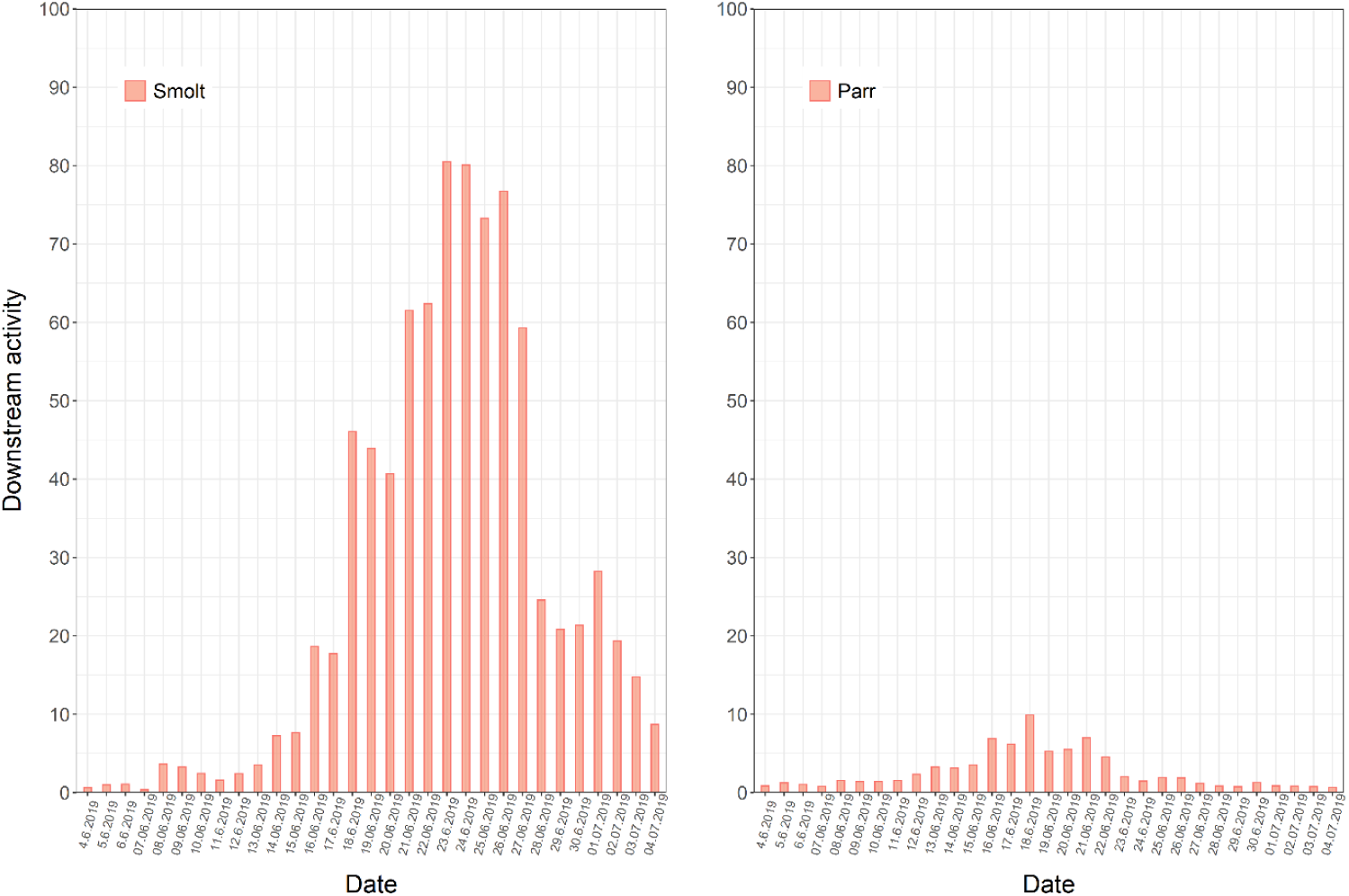
Best linear unbiased predictors (i.e. BLUPs) for downstream activity for each date. BLUPs are presented separately for migrant (i.e. smolt) and non-migrant (i.e. parr) fish. BLUPs for each date are extracted from models presented in Table 1. As the results were qualitatively the same across all three behaviours, we present here only the BLUPs for downstream activity.

**Supplementary Figure 2.**
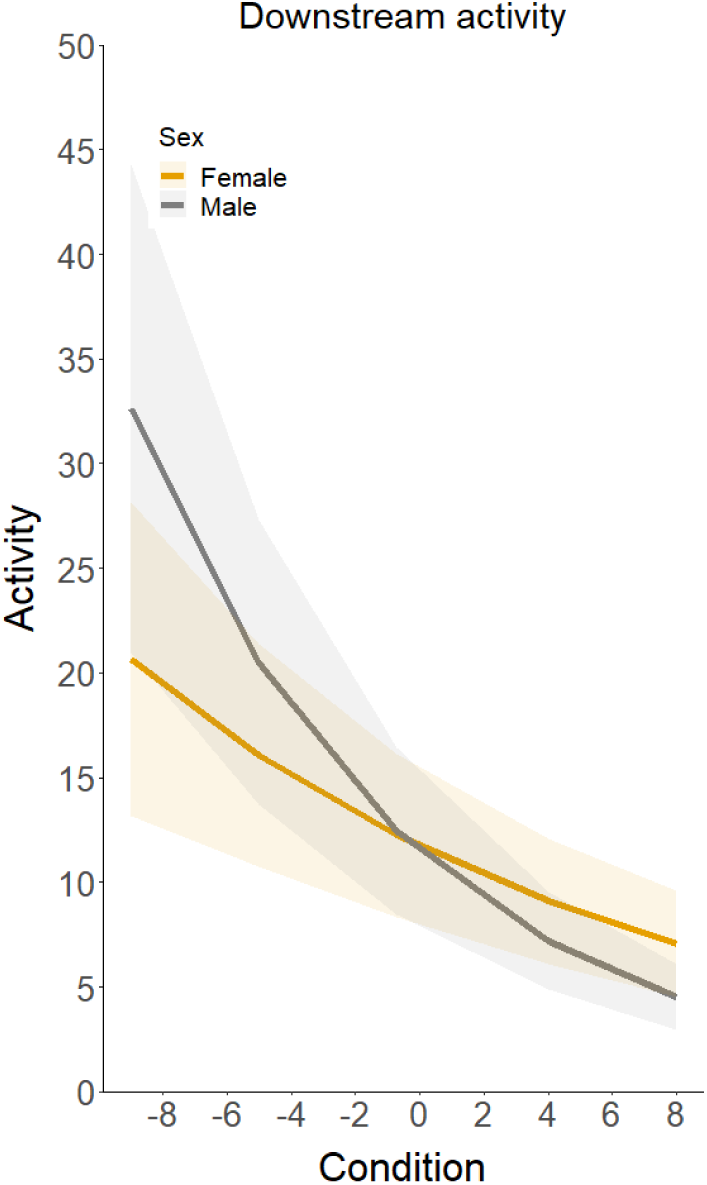
Linear predictions for sex-specific effects of body condition on recorded activity. Shaded areas represent standard errors around the predictions. The predictions are derived from model estimates presented in Supplementary Table 1. As the results were qualitatively the same across all three behaviours, we present here only the predictions for downstream activity. In the X-axis, negative values refer to lower condition while positive values refer to higher condition.

**Supplementary Figure 3.**
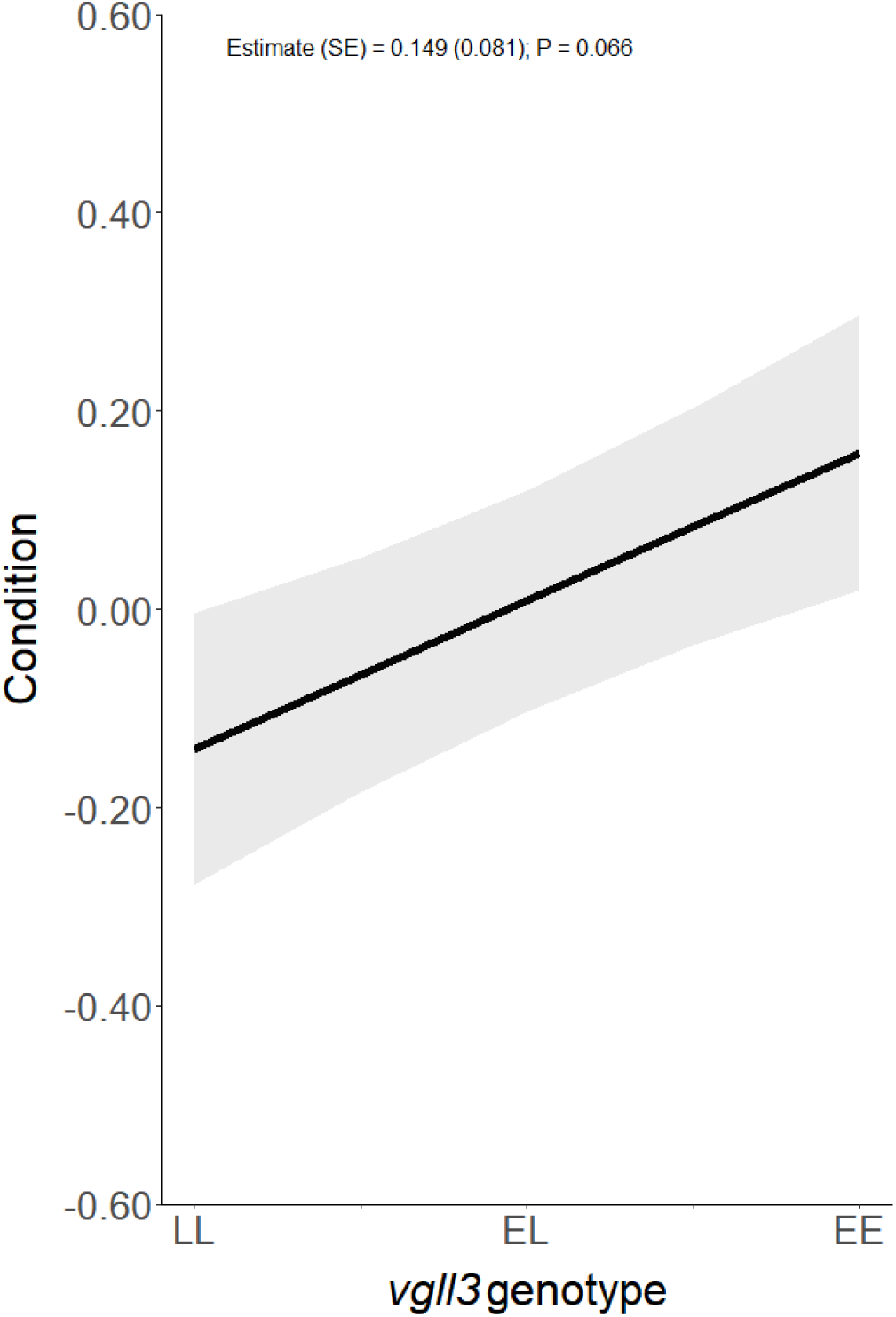
Linear prediction for the effect of *vgll3* genotype on condition among migrants. The shaded area represent standard errors around the prediction. The predictions are calculated from estimates delivered by a model: condition ∼ *vgll3*_additive + Sex + (1|Stream) + (1|mother) + (1|father). Other random effects, as present in the main models, were omitted since only one condition measurement per individual was obtained (thus among-individual, date and hour variation are not present). In the Y-axis, negative values refer to lower condition while positive values refer to higher condition.

